# Metal-Triggered Rheological Switching in Engineered Lanmodulin Condensates

**DOI:** 10.64898/2025.12.22.696036

**Authors:** Daniel S. Trettel, Brittany L. Huffman, Anna Clark, Xiaokun Yang, Babetta L. Marrone, C. Raul Gonzalez-Esquer

## Abstract

Reliable recovery of rare earth elements (REEs) is increasingly important for energy and communication technologies, yet current separation methods rely on harsh chemistries and are difficult to scale sustainably. Biological strategies offer a promising alternative but typically require immobilized proteins, introducing diffusion limitations and regeneration challenges. Here, we engineer a phase-separating variant of the high-affinity lanthanide-binding protein Lanmodulin (LanM) by fusing it to an encapsulation peptide (EP) microdomain. The resulting chimera, EP-LanM, undergoes liquid–liquid phase separation upon REE binding, forming condensates whose formation and dynamic properties are governed by metal stoichiometry and temperature. These condensates enable selective capture and release of La³⁺ and Nd³⁺ and support efficient recovery and reuse of the protein. Our findings show that metal-triggered protein condensation can be harnessed as an aqueous, reversible mechanism for REE enrichment, establishing a generalizable biophysical approach for designing functional phase-separating proteins for selective metal separation.

## Introduction

Advanced technologies increasingly depend on the reliable supply of critical metals that underpin energy, communication, and defense infrastructures.^1^ Among them, the rare earth elements (REEs) are indispensable components of high-performance magnets, displays, catalysts, and renewable-energy systems.^2^ However, REEs rarely occur in concentrated or pure mineral forms, requiring multistep chemical separations involving strong acids, organic solvents, and large energy inputs. These processes not only generate significant waste and, in some cases, hazardous by-products but also contribute to severe supply-chain and geopolitical vulnerabilities due to their concentration in a few global production centers.^3–5^ Developing sustainable, selective, and circular approaches for REE recovery is therefore an urgent technological and environmental priority.^6^

Biobased methods have recently emerged as promising alternatives to conventional chemical separation with reduced environmental impact.^7,8^ These methods stem from the increasing appreciation that diverse microorganisms have evolved ornate means of sequestering specific REEs to support biological processes.^9^ Among the most prominent examples is that of Lanmodulin (LanM), a protein chelator that exhibits exceptional selectivity for trivalent lanthanides through cooperative binding across four EF-hand motifs.^10,11^ LanM binds many REEs with picomolar affinity and can coordinate approximately three metal ions per molecule.^10^ Leveraging these properties, several groups have immobilized LanM on solid supports to perform affinity-based separations of REEs.^12–15^ While these systems demonstrate impressive specificity and recovery, their scalability is limited by diffusion and surface-access constraints and their reusability can be hindered by resin fouling and chemical regeneration requirements.^16–18^

Biomolecular condensation offers an underexplored, alternative paradigm for selective metal concentration.^19^ In such systems, biomolecules can spontaneously demix into dense and dilute phases, enabling the co-partitioning of bound ligands or cofactors.^20^ Phase separation thus acts as a dynamic, solution-phase enrichment process that is inherently scalable, dynamic, reversible, and free from immobilization steps. Despite the widespread biological use of phase separation for molecular organization, no naturally phase-separating REE-binding proteins have yet been identified.^21^

In this work, we bridge the gap between molecular coordination and mesoscale organization by engineering a phase-separating variant of LanM. By fusing the protein’s high-affinity binding domain to an encapsulation peptide (EP) microdomain,^22^ we created a system where liquid–liquid phase separation is strictly gated by REE binding. We demonstrate that the resulting condensates display tunable dynamic properties defined by metal stoichiometry, enabling a reversible, aqueous mechanism for selective REE enrichment. Ultimately, by establishing a direct link between ion occupancy and phase behavior, we provide a generalizable biophysical approach for converting specific molecular recognition into functional, macroscopic separation materials.

## Results

### Chimeric LanM phases separate in response to lanthanides

Lanmodulin (LanM) has not been previously observed to naturally undergo biomolecular condensation. As such, our first task was to append a condensation tag to LanM and observe its effects on protein behavior. We made two designs based on the *Methylorubrum extorquens* LanM with the native N-terminal signal peptide removed.^10^ In one case, the signal peptide was replaced by the native encapsulation peptide (EP) and linker from that of *Salmonella enterica* PduD (Figure 1A, Supplemental Table 1). This peptide was chosen for its small size, modularity, and noted ability to trigger liquid-liquid phase separation (LLPS) when used as a fusion molecule.^22^ This chimera (EP-LanM) was qualitatively assessed for its ability to undergo LLPS in response to specific buffer additives (Figure 1B). Interestingly, turbid solutions were only observed when both crowding agent and LaCl_3_ were present, with light microscopy confirming the presence of droplets. Meanwhile, previous work with EPs had only observed crowder and protein concentration effecting condensation.^22^

**Figure 1:**
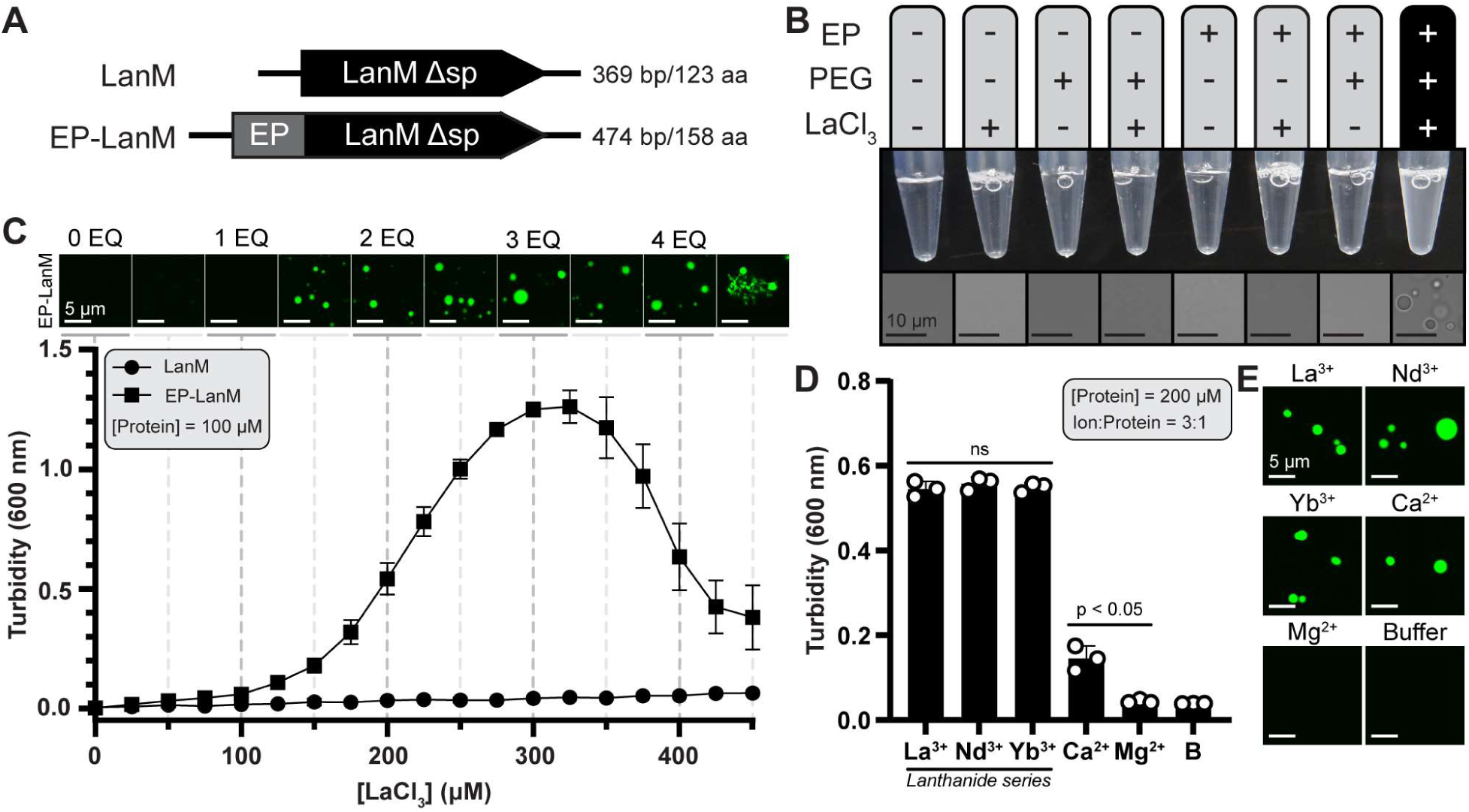
The EP microdomain supports Lanmodulin phase separation in the presence of known ligands. **(A)** A graphic schematic of the *M. extorquens* LanM constructs used in this study. Both lack the native signal peptide (Δsp). **(B)** A qualitative assessment of buffer additives that support phase separation was performed. Scale bars, 10 µm. **(C)** The turbidity response of LanM constructs as a function of LaCl_3_ concentration. Images of Alexa Fluor 488-labeled EP-LanM prepared at specified intervals are shown. The points on the curve represents the means ± standard deviation of three replicate experiments. Scale bars, 5 µm. **(D)** The turbidity response of EP-LanM to various ions was measured. No significant (ns) difference was determined among the lanthanides as given by an unpaired t-test. The bars represent the means ± standard deviations of three replicate experiments. **(E)** Images of samples in panel D prepared with fluorescently labeled EP-LanM. Scale bars, 5 µm.

We next sought to understand the dependence of EP-LanM condensation on LaCl_3_ concentration. For this, we paired turbidity measurements with laser scanning confocal microscopy of Alexa Fluor 488-labeled LanM and EP-LanM (Figure 1C, Supplemental Figure 1A). Turbidity is a common means of assessing LLPS based on light scattering caused by droplets in solution. While it should not be used to make quantitative statements about the number or size of droplets in solution,^23^ we found it agreed well with the total amount of condensed protein matter (Supplemental Figure 1B). This assay found that EP-LanM turbidity increased with LaCl_3_ until it reached a maximum at approximately 3 molar equivalents of metal added before it began to decline. Similarly, droplets were observed in all conditions after approximately 1 molar equivalent of metal addition, with aggregates present at high equivalents, perhaps explaining the decline in turbidity. LanM itself demonstrated no turbidity response and was not observed to form any structures in solution (Supplemental Figure 1A). With the metal dependence established, our investigation found that EP-LanM undergoes phase separation at concentrations as low as ∼10 µM in the presence of 3-molar equivalents of LaCl_3_ (Supplemental Figure 1C). Higher protein concentrations appeared to correlate with increased droplet size under these conditions (Supplemental Figure 1D).

Our initial screen suggests that turbidity, and therefore condensed material, peaks in the presence of 3 molar equivalents of metal. We used this to screen the metal specificity for LLPS of EP-LanM in the presence of La^3+^, Nd^3+^, Yb^3+^, Ca^2+^, Mg^2+^ (as chloride salts), and a buffer control. We chose these lanthanides based on their spread across the series, which can be important for binding, and broader technological appeal. Meanwhile, Ca^2+^ is a known weak binder to LanM^10^ and Mg^2+^ served as a negative control. Solutions of EP-LanM were prepared in the presence of 3-molar equivalents of metal and turbidity responses were measured. The highest turbidities were observed for lanthanides while Ca^2+^ showed a far weaker response (Figure 1D). No response was found for Mg^2+^ and buffer. The turbidity measurements were corroborated with microscopy imaging that confirmed the presence of droplets in response to La^3+^, Nd^3+^, Yb^3+^, and Ca^2+^ (Figure 1E). Altogether, this data demonstrates that EP-LanM will undergo LLPS in response to known ligands of the LanM domain.

### Stoichiometric metal binding drives EP-LanM condensation

Our initial data supports a relationship between metal stoichiometry and condensation. Specifically, condensation of EP-LanM peaked in the presence of 3-molar equivalents of LaCl_3_, which agrees with prior observations that LanM has three high-affinity binding sites.^11^ If binding site occupancy is related to protein condensation, then we reasonably expect that this relationship should hold at different concentrations of EP-LanM. To test this, we prepared EP-LanM at three different concentrations (75, 100, and 125 µM) and measured the turbidity in response to LaCl_3_ as before. As expected, the different EP-LanM concentrations led to different curve shifts and maxima, which could then be converted to molar equivalents of LaCl_3_ and turbidity normalized (Supplemental Figure 2). This revealed that the three curves nearly overlapped, with peaks close to 3-molar equivalents and similar half-maximal effective concentration (EC50) values when fit with a sigmoidal curve (Figure 2A, Supplemental Table 2). This data leads us to believe that EP-LanM condensation is driven by a stoichiometric dependence between the metal ligand and the protein.

**Figure 2:**
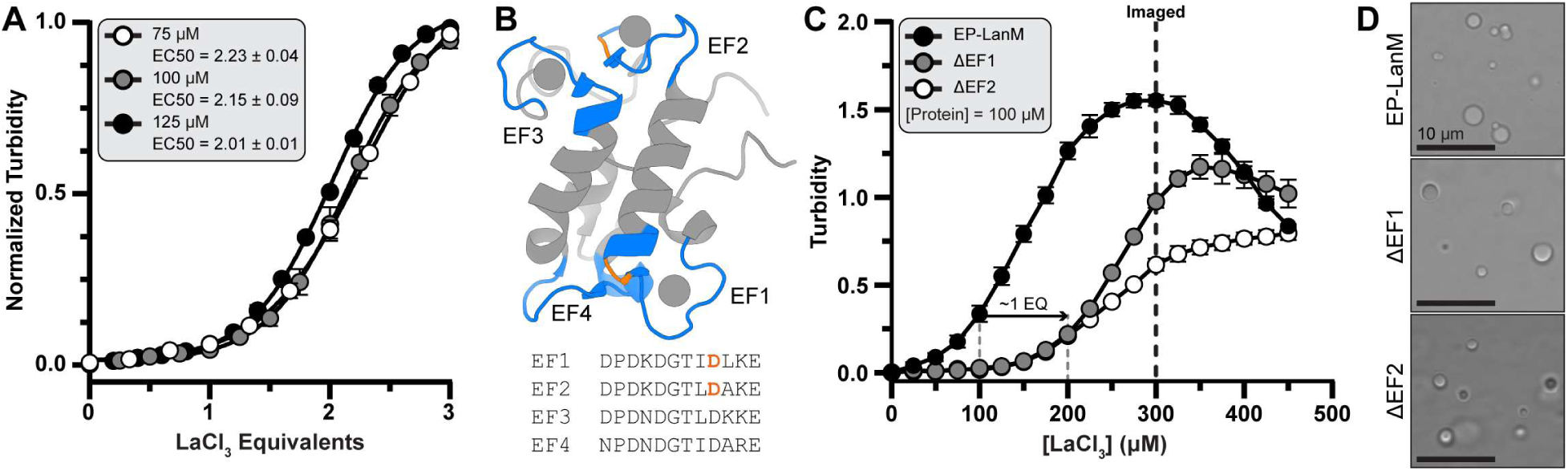
Chimeric Lanmodulin phase separation is related to ion binding capacity. **(A)** Turbidity curves of EP-LanM in response to LaCl_3_ were prepared at different protein concentrations. Their overlap suggests a stoichiometric binding effect. The data points are fit with a sigmoidal equation described in the Methods. **(B)** The structure of Y3+-bound LanM (PDB: 6MI5) is presented to show the four EF-hands (colored blue). Mutation sights are highlighted orange. **(C)** The turbidity curves of binding mutants are compared. Specifically, a D9A mutation was made to both EF1 and EF2. Three replicate experiments were performed individually for each sample in panels A and B and the means ± standard deviations are plotted. **(D)** Representative images of samples in panel B. Scale bars, 10 µm.

The observed stoichiometric dependence of the metal ligand does not explicitly tie ligand binding to being the driver behind EP-LanM condensation. To confirm that phase separation is strictly gated by ligand occupancy, we introduced D9A mutations to disable specific EF-hands (ΔEF1 and ΔEF2, Figure 2B).^24,25^ We hypothesized that if the stoichiometry of metal binding effected EP-LanM condensation, then we should see those effects displayed in their turbidity responses to LaCl_3_. Strikingly, the phase transition thresholds for these variants shifted by exactly one molar equivalent relative to the wild type (Figure 2C). This quantized shift indicates that the EP-LanM system functions as a molecular counter: the thermodynamic driving force for condensation is linearly coupled to the number of coordinated metal ions, allowing for precise stoichiometric tuning of the phase boundary. The different maxima may reflect different extents of conformational switching between ΔEF1 and ΔEF2. Regardless, all variants can produce droplets in solution (Figure 2D). This data demonstrates that EP-LanM condensation is driven by stoichiometric metal binding.

### Extent of metal binding tunes condensate dynamic properties

Our results so far demonstrate that the extent of metal binding can control the equilibria between the dense and soluble phases. Previous work has shown that metal stoichiometry also affects LanM conformational switching from a disordered state to a predominantly helical conformation.^10^ Taken together, these observations in which metal binding affecting both phase equilibria and conformational state suggest that the dynamical properties of EP-LanM condensates may be tuned by the extent of metal binding. As a baseline, we observed that Alexa Fluor 488-labeled EP-LanM samples demonstrated partial liquid behaviors such as droplet merging (Figure 3A).

**Figure 3:**
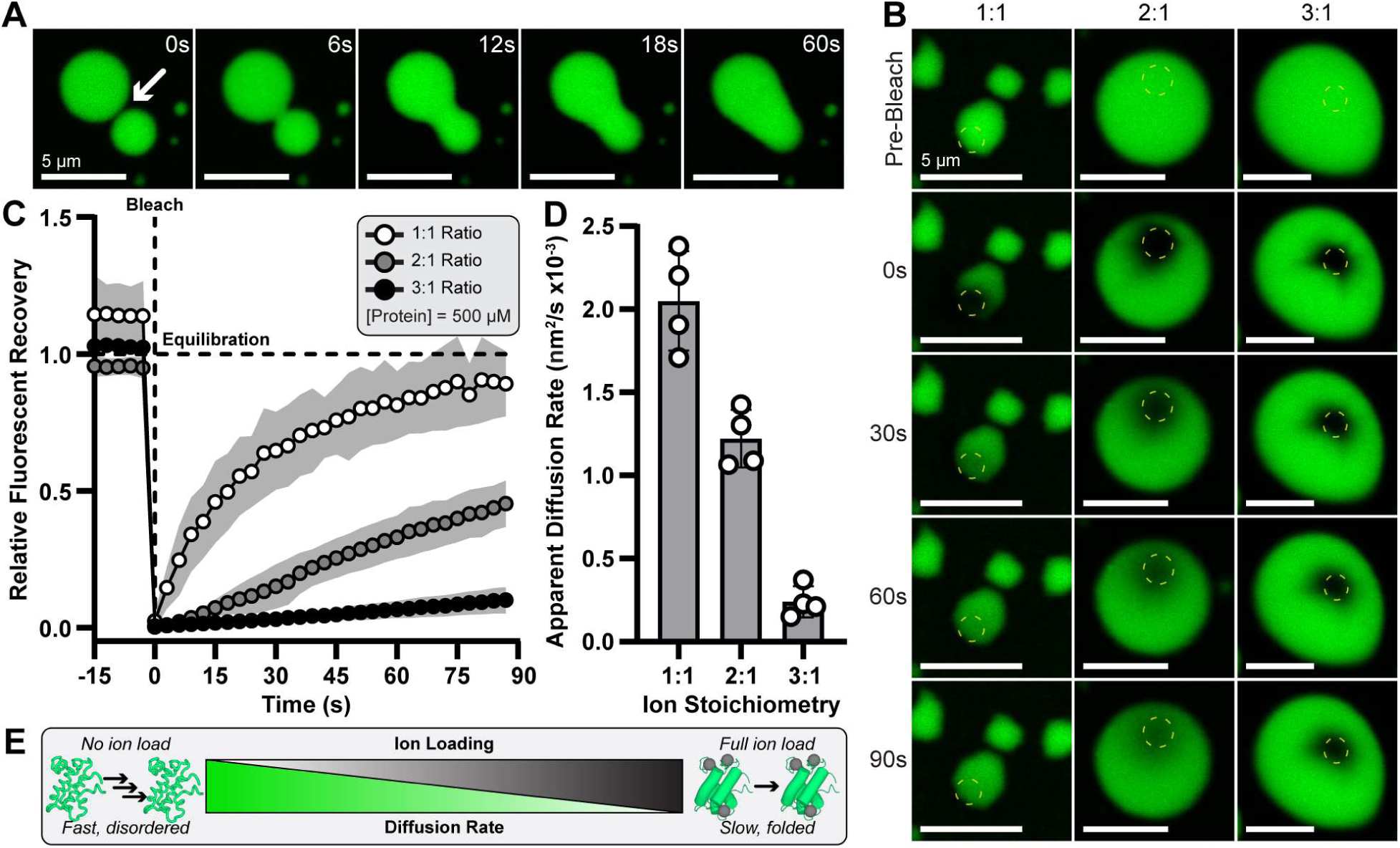
Chimeric Lanmodulin droplets are liquid-like with ion-tunable material properties. **(A)** Timelapse imaging of two fluorescently labeled EP-LanM droplets coming into contact and fusing. Samples were prepared at 250 µM protein at a 3:1 molar ratio of LaCl_3_. The arrow indicates the fusion point. Scale bars, 5 µm. **(B)** Fluorescent recovery after photobleaching (FRAP) experiments were performed on samples with different molar equivalents of LaCl_3_. Samples are shown before bleaching and at 30s intervals of their recovery. The yellow dashed circle represents the bleached space. Scale bars, 5 µm. **(C)** Quantification of FRAP with sampling every 3s demonstrates different recovery profiles. **(D)** FRAP profiles were used to calculate intra-droplet diffusion rates as a function of ion stoichiometry. Data in panels C and D is the means ± standard deviations of data collected from four separate droplets. **(E)** A rheological switch model is presented where protein diffusion and ion loading are inversely related.

Given our observations that metal binding affects phase equilibria and conformational switching, we wanted to understand if metal binding also affects condensate dynamical behavior. Towards this, we used the fluorescent recovery after photobleaching (FRAP) of droplets as a function of metal stoichiometry to probe molecular mobility.^26^ Here, we photobleached portions of labeled EP-LanM droplets prepared with different stoichiometries of LaCl_3_ and compared the recovery of that spot to an unbleached spot within the same droplet. The timescale of recovery, or intra-droplet equilibration, yields information about protein mobility within the condensates.^27,28^ Samples of EP-LanM at 500 µM with 1, 2, or 3 molar metal equivalents were prepared and FRAP was performed (Figure 3B). Images were taken in 3s intervals beginning 15s prior to bleaching and up to 90s afterwards (Figure 3C). The 1:1 condition showed a rapid, near-complete re-equilibration profile within this timeframe. The 2:1 condition showed slower recovery while the 3:1 showed the slowest recovery. The recovery times were extracted from each individual replicate and, in conjunction with the radius of bleaching, used to determine apparent diffusion coefficients (see Methods, Supplemental Table 3). The calculated apparent diffusion coefficients spanned nearly an order of magnitude and were, in increasing metal stoichiometry, 2050 ± 300, 1221 ± 174, and 241 ± 94 nm^2^/s (Figure 3D). This data indicates that metal addition leads to slower dynamics within EP-LanM condensates. We also varied the total protein concentration within samples and found no effect on the FRAP recovery profiles and apparent diffusion coefficients (Supplemental Figures 3C, 3D), suggesting that protein dynamics within the condensate are only dependent on the extent of metal binding. Unexpectedly, we observed that high metal stoichiometries (which induce LanM folding)^10^ resulted in a ten-fold decrease in intra-droplet diffusion coefficients. This inverse relationship suggests a ’rheological switch’ mechanism: while metal binding compacts the individual polypeptide, it simultaneously exposes motifs that enhance inter-chain networking. Consequently, the coordination of the third metal ion does not merely saturate the binding site but actively transitions the condensate from a fluid liquid to a dynamically arrested, high-viscosity network (Figure 3E).

### EP-LanM droplets are temperature responsive

We next sought to understand if simple environmental changes, like temperature, can control the phase equilibria of EP-LanM condensates. This could be particularly relevant in industrial settings to maximize REE capture. We tested the temperature dependence of condensation by cycling EP-LanM droplets, prepared at a 3:1 ratio, between 4°C, 22°C, and 37°C and measured the turbidity after equilibration (Figure 4A). The system displayed reversible, temperature-dependent demixing, with turbidity increasing at lower temperatures. This profile is consistent with enthalpically driven phase separation, where the favorable energetic gain of protein-protein/protein-metal interactions in the dense phase overcomes the entropic penalty of demixing. This thermal reversibility confirms that the condensates remain in thermodynamic equilibrium, allowing temperature to be used as a non-chemical switch for capture and release.

**Figure 4:**
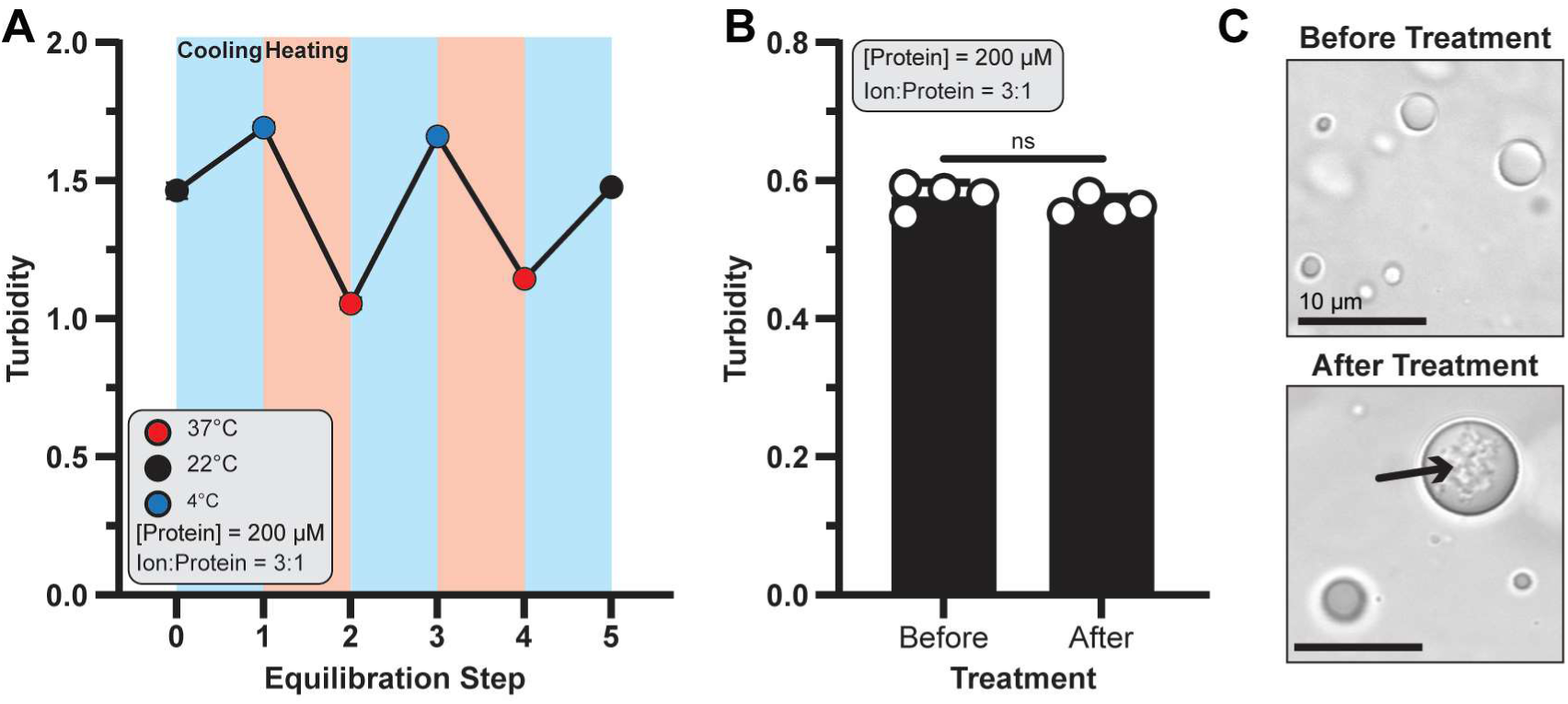
Chimeric Lanmodulin droplets are temperature responsive and reversible. **(A)** Turbidity of EP-LanM samples was measured in response to temperature changes. Dot color represents the temperature of the sample at measurement, while background color represents whether the sample was undergoing heating or cooling between measurements. Data are the means ± standard deviations of three replicate experiments. **(B)** Turbidity was measured before and after heat treatment at 95°C. No significant (ns) difference was determined by an unpaired t-test. Data are the means ± standard deviations of four replicate experiments. **(C)** Representative images of samples in panel B demonstrate droplet re-formation, albeit with internalized aggregates (arrow). Scale bars, 10 µm.

We next sought to understand the temperature limits of EP-LanM. Previous work has demonstrated that LanM-REE complexes can withstand temperatures upwards of 95°C,^29^ but it remains unclear if the EP microdomain can too. We tested this by measuring the turbidity of condensed EP-LanM samples before and after heat treatment at 95°C. Treated samples became visually clear at 95°C (not shown) but regained near-equivalent turbidity to their starting condition upon cooling back to room temperature (Figure 4B). Sample imaging showed that droplets reformed after heat treatment, albeit with morphological distinctions (Figure 4C). Specifically, the heat-treated samples displayed larger droplets enveloping seemingly aggregated inclusions which were observed in all four replicates (Supplemental Figure 4A). SDS-PAGE analysis of samples prepared in the presence or absence of LaCl_3_ demonstrated similar banding patterns regardless of heat treatment (Supplemental Figure 4B). Interestingly, samples prepared with LaCl_3_ showed an electrophoretic shift towards higher masses. We found that this shift was induced by LaCl_3_ addition and likely suggests that LanM-REE complexes can survive SDS-PAGE sample preparation, preserving their conformational switching which is reflected in their electrophoretic mobility (Supplemental Figure 4C). This data also rules out metal-driven protein hydrolysis as a source of the aggregates. We believe these structures are likely to be dynamically arrested, out-of-equilibrium states resulting from the heating and cooling of condensates with slow internal dynamics.^30,31^ Additionally, some fraction of EP microdomains remain partially disrupted during cooling, leading to general aggregation which then act as nucleation sites. Regardless, the EP microdomain is robust enough to still support high levels of protein condensation without any sequence optimization.

### Protein condensation supports high levels of metal and protein recovery

Our data points towards two major practical considerations towards using the EP-LanM system for metal extraction; (1) the molar ratio of metal and protein and (2) the temperature at which condensation occurs. These are both crucial as they control the extent of condensation and therefore the amount of captured material. With this in mind, we developed a bench-scale protocol for REE extraction using protein phase separation (Figure 5A). Here, the condensation reaction was mixed and equilibrated on ice to shift equilibrium towards the dense phase. The droplets were then enriched via centrifugation and resuspended in a low pH buffer. This buffer stimulates metal release and therefore EP-LanM solubilization. The solubilized metal and proteins were then separated by mass using a filtration unit. The metal and protein fractions were then independently quantified with inductively coupled plasma atomic emission spectroscopy (ICP-AES) and A_280_ measurements, respectively. We tested three conditions using this workflow: (1) a simple extraction of La^3+^, (2) a complex extraction of La^3+^ in the presence of excess Ca^2+^ (5x over lanthanum), and (3) a co-extraction of La^3+^ and Nd^3+^ (Figure 5B, conditions 1-3, respectively).

**Figure 5:**
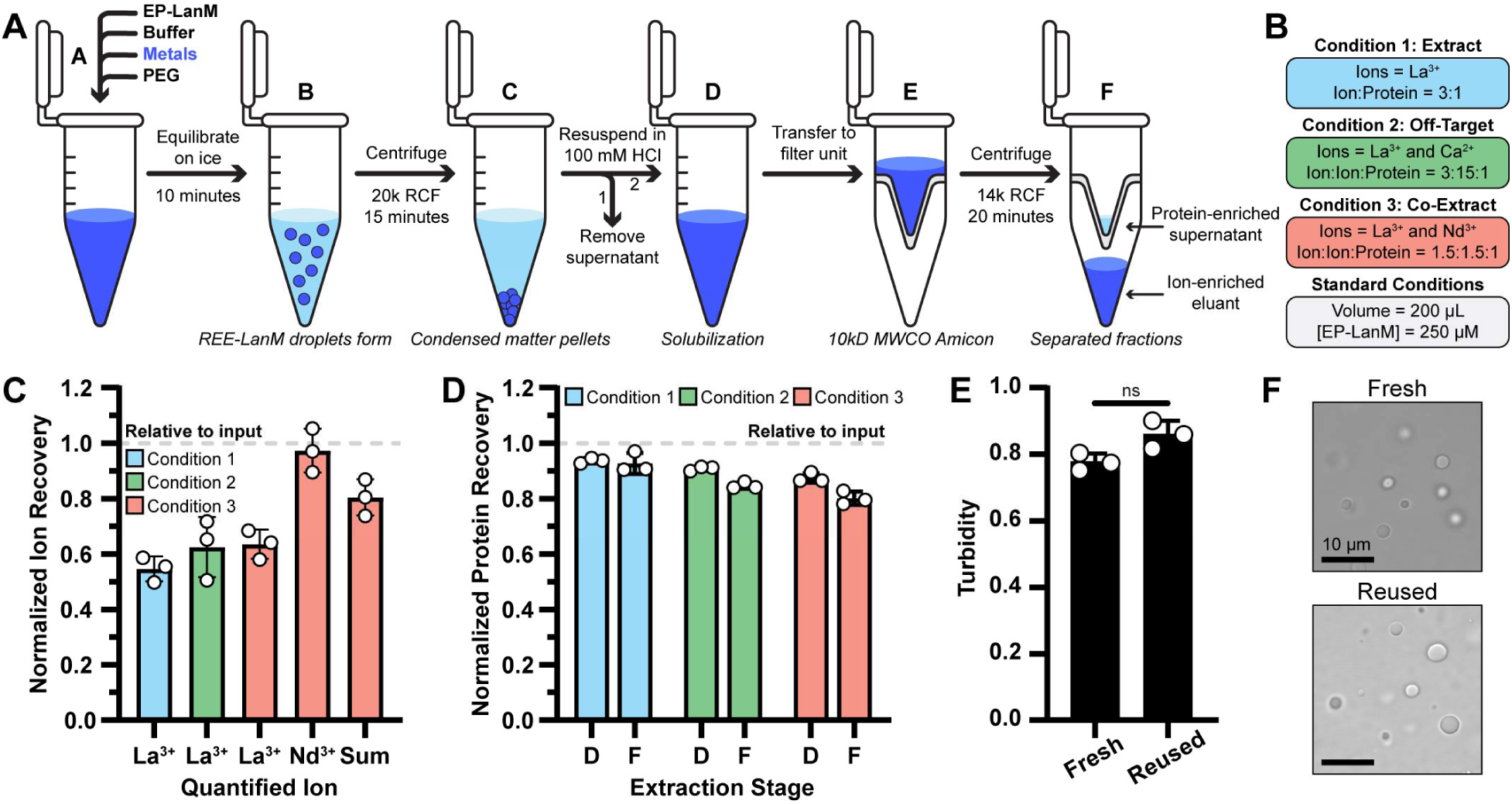
Leveraging protein phase separation for elemental extraction. **(A)** A workflow for a bench-scale ion extraction assay is shown. Here, components are mixed and equilibrated on ice. Condensates are captured via pelleting prior to ion release stimulated by pH drop. The extract is then filtered to separate protein from extracted ions. **(B)** Three extraction conditions were tested and differed by their ion composition. **(C)** The ion-enriched eluant was measured with ICP-MS and percent recovery was determined relative to the initial inputs. **(D)** Protein recovery for each condition was measured after steps D (condensate resuspension) and F (final fraction). **(E)** Reusability is assessed by comparing the turbidity of freshly prepared samples to samples previously used for a round of extraction. No significant (ns) difference was found between samples as determined by a Welch’s t-test. Three replicate experiments were performed individually for each sample in panels C-E and the means ± standard deviations are plotted. **(F)** Images of fresh and reused samples from panel E are shown with similar morphology. Scale bars, 10 µm.

Metal-enriched eluants were analyzed with ICP-AES and the capture efficiency was determined relative to the input values (Figure 5C). Conditions 1-3 extracted 54.6% ± 4.5%, 62.5% ± 10.9%, and 63.5% ± 5.3% of expected La^3+^, respectively. Condition 3 also extracted 97.3% ± 7.9% of expected Nd^3+^, leading to a combined extraction rate for both La^3+^ and Nd^3+^ (labeled ‘sum’ in Figure 5C) of 80.4% ± 6.5%. Carry over sodium, used in the condensation buffer (50 mM input in all samples), and Ca^2+^ (3750 µM in Condition 2) were detected but at levels between the limits of quantification (LOQ) and detection (LOD), placing upper and lower bounds on the extent of Na^+^ and Ca^2+^ contamination. This allowed us to estimate purification factor ranges for La^3+^ over Na^+^ (2.1-6.8) or Ca^2+^ (6.5-21.8) and Nd^3+^ over Na^+^ (3.6-11.9), confirming high specificity (Supplemental Figure 5). Protein recovery was also assessed at two stages of the workflow, either after condensate pelleting or at the final fraction step (Figure 5A, labeled steps D and F). All samples demonstrated high levels of protein recovery with minimal losses between condensate pelleting and filtration (Figure 5D). Final protein recovery for conditions 1-3 were 92.7% ± 3.8%, 84.7% ± 1.2%, and 80.1% ± 2.4%, respectively. These results demonstrate that protein phase separation can be used to efficiently extract REE’s and recover the input protein.

While input protein can be recovered at high levels, it remained unclear if the recovered protein can still be responsive to metals and therefore be reused. To test this, we compared the ability of fresh or previously used EP-LanM to condense in the presence of LaCl_3_ (Figure 5E). We found that reused EP-LanM can achieve similar levels of condensation as fresh EP-LanM. Further, microscopy imaging revealed that droplets between the two samples were indistinguishable from one another (Figure 5F). This data shows that the extraction process has minimal to no effect on subsequent EP-LanM condensation and that, in principle, these droplets may be able to support subsequent rounds of extraction.

## Discussion

The ability to translate precise molecular recognition events into macroscopic phase transitions represents a new frontier in the design of responsive biomaterials. While natural biomolecular condensates dynamically respond to cellular cues, rationally coupling high-affinity metal coordination to phase separation offers a unique mechanism to program mesoscale material properties. Here, we demonstrate that the conformational dynamics of a lanthanide-binding protein can be harnessed to drive liquid–liquid phase separation, establishing a direct link between ligand stoichiometry and droplet viscosity. By engineering a system where metal binding inversely regulates bulk diffusion, we show that the thermodynamics of specific ion coordination can be effectively transduced into reversible phase equilibria, providing a generalizable biophysical approach for aqueous metal separations.

Our implementation of the EP-LanM protein demonstrates distinct advantages over immobilization-based strategies. First, EP-LanM leverages conformational switching to support condensation at high metal stoichiometries. As a biophysical phenomenon, this entails inherent scalability and reversible control via simple changes to temperature, pH, and concentration. Crucially, avoiding solid supports eliminates tedious immobilization steps that can limit the conformational freedom disordered proteins like LanM require.^32^ Unlike beads which are prone to fouling, diffusion limitations, and thermal instability,^16–18^ our aqueous approach survives high temperatures and maintains high surface access. EP-LanM also demonstrated high purifications factors over Na⁺ and Ca²⁺ that can likely be further enhanced with addition of standard wash steps. Additionally, successful reuse suggests system longevity is limited only by intrinsic protein stability. Our work with full-length LanM also differs from aggregation by lanthanide-binding tags.^33^ By using native-like LanM, we retain the exquisitely high affinity and selectively while building in a dynamic organizational component that regulates phase behavior.

Crucially, our data establishes a direct, inverse relationship between the extent of metal coordination and bulk protein diffusion. We observed that increasing metal stoichiometry results in a progressive reduction in molecular rearrangements within the droplet. This seemingly presents a biophysical paradox: while metal binding triggers the folding of LanM into a compact, helical state, which theoretically reduces the hydrodynamic radius and should accelerate diffusion, it simultaneously arrests bulk internal dynamics. We propose that this rheological switch arises because the metal-stabilized conformation exposes specific interaction motifs that drive inter-molecular networking rather than simple packing. Consequently, the coordination of the third metal ion acts as a critical gate, transitioning the condensate from a fluid liquid to a dynamically arrested, high-viscosity network we estimate to be ∼500 Pa·s (see Methods). We note that this is a rough estimate, as Stokes-Einstein assumptions commonly break in dense condensates and that further insight is needed from single-molecule tracking, microrheology, and binding kinetics experimentation. Still, this suggests that LanM’s conformational switching regulates the system’s effective connectivity, converting thermodynamic binding energy into macroscopic mechanical stability.

The synergistic coupling of the EP microdomain’s self-association with LanM’s conformational switching provides a modular framework for rationally engineering phase behavior. By linking the driver of phase separation (the EP tag) to a conformational trigger (metal binding), we establish a design space where the saturation concentration and ligand specificity can be tuned independently. For instance, increasing the system’s valency, either through tandem duplication of the EP domain^22^ or by employing LanM variants that obligately dimerize upon chelation,^13^ could promote condensation at lower protein concentrations and enhance ion partitioning. Furthermore, because the EP microdomain only supports LLPS when coupled to the holo-state of LanM, the emergent protein-protein interfaces within the dense phase represent a new, computationally accessible axis for optimization. This logic suggests that future designs can move beyond simple affinity tags to create ’smart’ extraction fluids, where the quaternary topology of the fusion construct is programmed to maximize recovery efficiency, minimize background salt sensitivity, and lower crowder requirements.

Altogether, our findings demonstrate that liquid–liquid phase separation can be rationally engineered into a high-affinity metalloprotein to create a responsive and reusable platform for rare-earth extraction. EP-LanM couples precise lanthanide recognition with metal-dependent condensate formation, enabling stoichiometry-tuned rheological properties, efficient ion capture, and near-quantitative protein recovery without reliance on immobilization or harsh chemical processing. By integrating molecular recognition with emergent mesoscale organization, this work establishes protein condensation as a generalizable biophysical strategy for selective metal enrichment and highlights new opportunities to design functional phase-separating proteins for broader separation and sustainability applications.

## Methods

### Molecular Biology

The Lanmodulin (LanM) gene was sourced from *M. extorquens* with N-terminal signal peptide (first 21 amino acids) removed. Variants of this were designed by appending the *S. enterica* PduD encapsulation peptide sequence and its native linker to the N-terminus of LanM (EP-LanM) and knockout designs were made to EF1 and EF2 by single alanine substitutions. All designs included a C-terminal hexahistidine tag appended via a GGS linker. These designs were ordered from Twist Biosciences as synthetic gene fragments codon optimized for *E. coli*. Gene fragments were cloned via Gibson assembly strategy into a pET11a backbone PCR linearized with primers oDT122 and oDT123. The PCR product was treated with DpnI (NEB) restriction enzyme prior to purification with a Zymo DNA Clean and Concentrator kit. Gibson assembly was performed using NEB HiFi assembly mix using a 3:1 molar ratio of insert to backbone for 20 minutes at 50°C. Assemblies were frozen at -20°C prior to transformation into NEB5α competent cells. Correct clones were identified by whole-plasmid sequencing provided by Plasmidsaurus.

### Protein Purification

All proteins purified used the following protocol. Plasmid DNA was transformed fresh into NEB T7 Express cells and plated onto LB plates supplemented with carbenicillin at 100 µg/mL (this supplementation is consistent hereafter) and grown overnight at 37°C. The next day, a single colony was used to inoculate 5 mL of 2xYT media and grown overnight at 37°C, 250 RPM. The following day, this starter culture was used to seed 500 mL of 2xYT media in a 2 L baffled flask and grown until mid-log (OD600 = 0.60 – 0.80) where the cultures were then briefly cooled to room temperature prior to addition of IPTG to 100 µM to induce protein expression. The flask was then returned to the incubator to incubate for 16-18 hours at 22°C, 250 RPM. The next morning, the culture was pelleted at 5000 RCF for 15 minutes. The supernatant media was removed, and the pellet transferred to a 50 mL conical tube and frozen at -20°C for at least 30 minutes. For lysis, the pellet was thawed and resuspended in NP-40 High Salt Lysis Buffer (3 mL/g, RPI) supplemented with 12.5 U/mL benzonase (Syd Labs), 0.5 mg/mL egg white lysozyme, 5 mM MgCl_2_, and 0.5 mM PMSF. The slurry was incubated for 30-60 minutes in an incubator set to 30°C, 250 RPM. The insoluble matter was then removed by centrifugation at 16000 RCF at 4°C for 15 minutes. The supernatant was then mixed with 1 mL of PureCube 100 Ni-INDIGO Agarose beads pre-equilibrated into Buffer A (10 mM Tris-HCl, 50 mM NaCl pH 8.0) and incubated in an end-over-end mixer for 30 minutes at room temperature. Purification then proceeded by gravity flow. After the flowthrough was drained, the beads were washed in 3 sets of 5 column-volumes of Buffer B (Buffer A + 25 mM imidazole) and protein was eluted in 3 sets of 1 column-volumes of Buffer C (Buffer A + 400 mM imidazole). Eluted protein was then dialyzed against 2x1 L batches of Buffer A in a 10 kD MWCO Slide-A-Lyzer for a minimum of 8 hours each at 4°C with stirring. Final protein concentration was determined via Nanodrop measurement at 275 nm using an extinction coefficient of 1400 M^-1^cm^-1.10^

### Fluorescent Conjugation

LanM and EP-LanM were diluted to 1 mM in 500 µL from concentration stocks and dialyzed into 0.1 M sodium bicarbonate pH 8.5. For each labeling reaction, 100 µg of Alexa Fluor 488 TFP ester (Thermo Fisher) was resuspended in 10 µL of anhydrous DMSO and immediately added to the protein solution. The protein-dye solution was vortexed for 5 seconds then incubated in a shaking incubator set to 25°C for 1 hour. The remaining dye was separated with a gravity PD10 column equilibrated into Buffer A. The first fluorescent band was captured as an approximately 1 mL fraction containing labeled protein. The eluant was quantified by measurement on a Nanodrop. Further details on protein quantification and dye:protein ratio can be found in Supplemental Table 4. In the cases where fluorescently LanM derivates were used, fluorescently labeled protein was added to account for 10% of the final concentration. Given the labeling efficiencies described in Supplemental Table 4, this meant labeled protein accounted for <2% of each the final concentration leading to minimal perturbation.

### Turbidity Assays

All turbidity assays were performed in Condensation Buffer (10 mM Tris-HCl, 50 mM NaCl, 25% (w/v) PEG2K pH 8.0) and all turbidity measurements were performed at 600 nm. In all cases, turbidity was triggered by adding a 2x solution (50% w/v) of PEG2K last. Effects of La^3+^ equivalents: Assays measuring the condensation of mutant LanM as a function of La^3+^ concentration occurred at room temperature in 1 mL triplicate volumes at 100 µM protein with LaCl_3_ added serially 1 µL per measurement from a 25 mM stock solution. This was chosen such that each addition corresponded to 0.25 molar equivalents of La^3+^ addition and the summed addition equaled <2% the total volume, making volume changes negligible. For each measurement, LaCl_3_ was added followed by pipet mixing and equilibration for 30 seconds prior to measurement in a cuvette reader.

### Minimal condensation concentration

This assay was performed via an automated Integra Assist Plus liquid handling system. Briefly, a master stock of LanM protein at 400 µM was prepared in a 3:1 molar ratio with LaCl_3_ in 10 mM Tris-HCl, 50 mM NaCl pH 8.0. This stock was then serially diluted in a half-area 96-well plate in quadruplicate before then being mixed with an equal volume of 10 mM Tris-HCl, 50 mM NaCl, 50% (w/v) PEG2K such that the final PEG2K concentration was 25%. The plate was then incubated at room temperature for 15 minutes before measurement.

### Pelleting assay

Triplicate samples of LanM and EP-LanM (with Alexa Fluor labeled spike-in) were prepared in Condensation Buffer at 100 µM at 1 mL scale with the indicated ratios of LaCl_3_ at room temperature. After equilibration for 10 minutes, turbidities were measured in cuvettes then 200 µL was transferred to 1.5 mL microcentrifuge tubes and subjected to centrifugation at 20000 RCF for 5 minutes. After, the supernatants were removed and the pellets were resuspended. 50 µL fractions of each pellet and supernatant had their fluorescence quantified on a Biotek Synergy plate reader in 96-well black plates.

### Temperature Shift Assays

The temperature shift assay was performed in triplicate 96-well plate format. LanM variants were prepared at 200 µM with a 3:1 ratio of LaCl_3_ in Condensation Buffer at 100 µL total volume. The mixtures were incubated at each step for 15 minutes prior to measurement and incubation at the next temperature. In the heat treatment assay, samples again were prepared at 200 µM protein at a 3:1 ratio of LaCl_3_ in Condensation Buffer in 200 µL total volume in quadruplicate PCR tubes. Each sample was split in half with one set receiving treatment and the other left at room temperature. The treatment set at 95°C for 4 minutes and slowly cooled to 25°C over 75 minutes. 50 µL of each sample was then measured for turbidity in a half-area 96-well plate in a Biotek Synergy plate reader.

### Microscope Imaging

All imaging was performed on an Olympus FV3000 confocal microscope using an 100x objective lens with immersion oil. All samples were imaged by aliquoting 3-5 µL of solution onto a coverslip positioned on the stage prior to sandwiching the sample with another cover slip. All samples were imaged with transmitted light and, in the case of fluorescently labeled samples, using the 488-laser line for Alexa Fluor 488 excitation. All images are presented without further modifications beyond false-coloring (for fluorescent images) and cropping.

### Photobleaching Experiments

Photobleaching was performed at the indicated protein and ion concentrations, mixed at 200 µL final volume in Condensation Buffer. Due to the mobility of free droplets, photobleaching was performed on droplets that were adhered to the coverslip surface. Imaging was performed in 3 second intervals ranging from 15 seconds prior to bleaching to 90 seconds after the bleaching event. Bleaching itself was performed for 0.5 seconds at 50% intensity for the 488 laser. Image series were exported and quantified in ImageJ. Recovery was quantified and normalized as the ratio of the intensity of the background-subtracted bleached spot over the intensity of a background-subtracted unbleached spot within the same droplet. Recovery profiles were plotted and fit to the following equation in GraphPad Prism version 10.6.1:

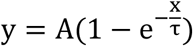

Where y is the normalized intensity versus time (x). A is the fraction of mobile proteins (set to 1, assumes all proteins are mobile across large time scales) and τ is the recovery half time in seconds.^34^ Diffusion rates were then calculated by dividing the square of the photobleached spot radius by the recovery time (D = 0.25r^2^/τ).^35^ Fit parameters, spot diameters, and diffusion rates for all individual replicates are listed in Supplemental Table 3. To estimate viscosity, we used a structure of LanM bound to 3 metal ions^11^ to estimate a Strokes radius of ∼2 nm with the Hullrad server^36^ and, given our calculated diffusion rates at the 3:1 ratio, estimate bulk apparent viscosities of ∼500 Pa·s using the Stokes-Einstein equation. We note that this is a rough estimation, as Stokes-Einstein assumptions commonly break in dense condensates.

### Mock Extraction Assay

Elemental extraction was performed in 1.5 mL microcentrifuge tubes with 250 µM protein in Condensation Buffer at 200 µL scale. Ions were added at either (1) 750 µM LaCl_3_, (2) 750 uM LaCl_3_, 3750 uM CaCl_2_, or (3) 375 uM LaCl_3_ and NdCl_3_. Condensation was triggered in a pre-massed tube by addition of PEG2K and samples were equilibrated at 4°C for 15 minutes. Droplets were then collected by centrifugation at 20000 RCF for 10 minutes at 4°C. The supernatant was carefully removed and transferred to another massed tube and the clear pellet was resuspended in 200 µL of 100 mM HCl. The pellet and supernatant tubes were then massed and protein contents quantified by Nanodrop. The pellet fraction was then filtered once through a 0.5 mL 10 kD MWCO Amicon filter at 14000 RCF for 30 minutes at 4°C. The unfiltered supernatant was collected into a massed tube and protein contents quantified. The eluant (pellet fraction filtrate) was similarly massed and stored for ICP-AES quantification.

#### ICP-AES

Concentrations of La^3+^ and Nd^3+^ were determined by inductively coupled plasma-atomic emission spectroscopy (ICP-AES) using a Perkin Elmer Optima 8000 instrument. Nd^3+^ and La^3+^ calibration standards were prepared from a series of stock solutions (3000 µM,1000 µM, 333 µM, 111 µM, and 37 µM Nd or La) serially diluted from 1 M stocks in 0.1 M HCl. A 150 µL aliquot from each stock solution was diluted to 2 mL in 0.1 M HCl, to create a five-point calibration curve (31.3 ppm, 10.4 ppm, 3.47 ppm, 1.16 ppm, and 0.39 ppm Nd or La). Multiple wavelengths were monitored to quantify La^3+^ and Nd^3+^. La^3+^ concentrations were quantified at 384.9 nm, 407.7 nm, and 408.7 nm. Nd^3+^ concentrations were quantified at 401.22 nm, 406.1 nm, 424.73 nm, and 430.35. A 150 µL aliquot from each sample was diluted to 2 mL in 0.1 M HCl for ICP-AES analysis. The average La^3+^ and Nd^3+^ concentrations for each sample were converted to µM and corrected for the dilution factor. Percent recovery was determined using the equation below where M_f_ is the final concentration of La^3+^ or Nd^3+^ and M_i_ is the initial concentration of La^3+^ or Nd^3+^.

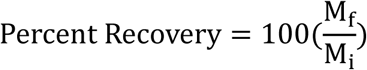

### Limit of quantification and limit of detection

The Limit of Quantification (LOQ) for La^3+^ and Nd^3+^ was determined from the calibration curve for each wavelength measured.^37^ This was performed using:

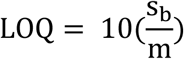

where s_b_ was the error in the y-intercept for the calibration curve and m was the slope of the calibration curve. Similarly, the Limit of Detection (LOD) was determined using:

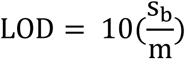

where s_b_ was the error in the y-intercept for the calibration curve and m was the slope of the calibration curve. Using these values, purification factors were calculated using the formula:

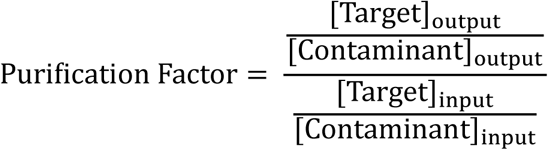

Where the targets were La^3+^ or Nd^3+^ and the contaminants were Na^+^ or Ca^2+^.

## Data Availability

All plasmids, strains, and data generated in this study are available by contacting the corresponding author. Source data are also provided with this paper.

## Acknowledgements

The authors would like to thank Dr. Ibraheem Alshareedah for providing thoughtful comments on this manuscript. The authors gratefully acknowledge the Laboratory Directed Research and Development (LDRD) project no 20240001DR and the U.S. Department of Energy through the Los Alamos National Laboratory. Los Alamos National Laboratory is operated by Triad National Security, LLC. For the National Nuclear Security Administration of the U.S. Department of Energy (contract no. 89233218CNA000001).

## Author Information

### Authors and Affiliations

Biosciences Division, Microbial and Biome Sciences Group, Los Alamos National Laboratory, Los Alamos, NM, USA

Daniel S. Trettel, Anna Clark, Babetta L. Marrone, C. Raul Gonzalez-Esquer Plutonium Supply & Disposition Division, Los Alamos National Laboratory, Los Alamos, NM, USA

Brittany Leigh Huffman Chemistry Division, Inorganic, Isotope, and Actinide Chemistry Group, Los Alamos National Laboratory, Los Alamos, NM, USA Xiaokun Yang

### Contributions

D.S.T and C.R.G.E conceptualized the project. D.S.T. designed the protein variants and carried out purification and *in vitro* characterization. D.S.T. performed fluorescent conjugation and all confocal microscopy imaging and data analysis. B.H. performed ICP-AES and subsequent analysis. A.C. helped with protein purification and *in vitro* characterization. D.S.T. wrote the paper with input from all authors.

### Corresponding Author

Correspondence to C. Raul Gonzalez-Esquer

## Ethics Declaration

The authors have no conflicts of interest to declare.

