## Supplemental Information for "Metal-Triggered Rheological Switching in Engineered Lanmodulin Condensates"

### Engineered Protein Condensates Enable Aqueous Lanthanide Extraction

Supplemental Information

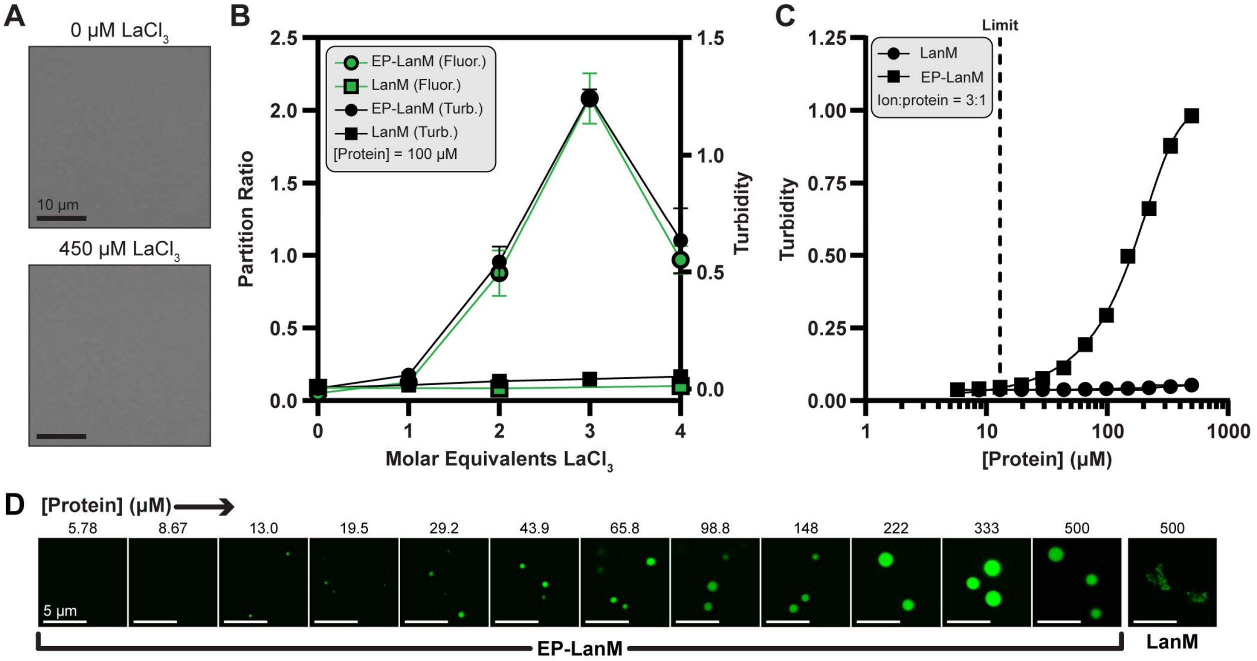


**<u>Supplemental Figure 1:</u> Materials related to main Figure 1. (A)** Light microscopy of LanM with either 0 or 450 μM $LaCl_3$ found no observable structures in solution. Scale bars are 10 μm. **(B)** Turbidity data is overlayed with the partition ratio. The partition ratio is the ratio of fluorescence in the dense phase over the dilute phase when turbid samples are pelleted. The mean ± the standard deviation of 3 experimental replicates is plotted. **(C)** The effect of protein concentration was determined by turbidity assay. Master stocks of LanM and EP-LanM were prepared in 3x molar excess of $LaCl_3$ and serially diluted before PEG2K addition. The means ± standard deviations of 4 experimental replicates are plotted. **(D)** Laser scanning confocal images from Supplemental Figure 3C are presented. The onset of droplets was observed at a minimum of 13.0 μM EP-LanM (labeled limit). Images are representative of a single replicate. All scale bars are 5 μm.

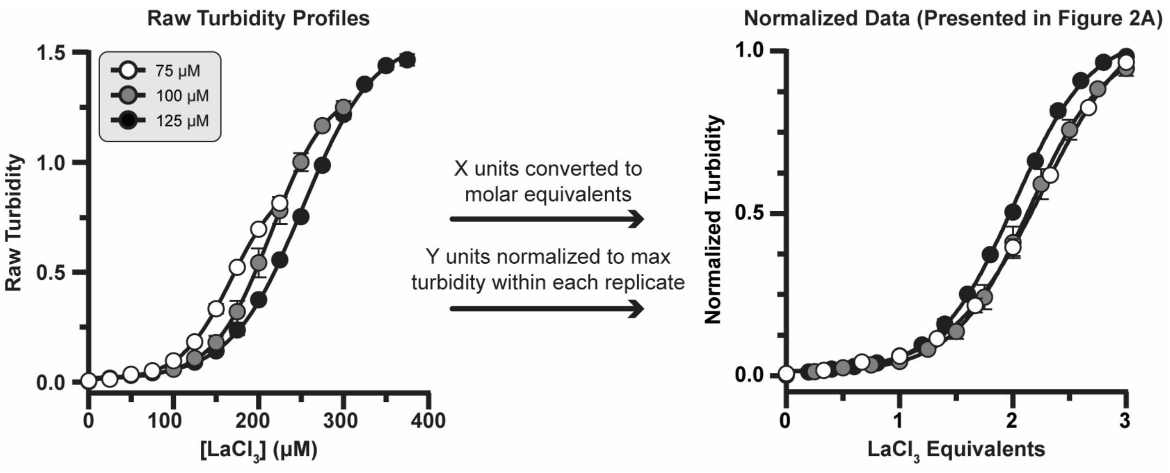


**<u>Supplemental Figure 2:</u> Materials related to main Figure 2.** The raw curves of data presented in Figure 2A are presented on the left. As these were performed at different protein concentrations, their maxima and x-axis shifts differed. The data was transformed by converting the x axis units to molar equivalents of $LaCl_3$ and the y axis had each replicate normalized by the max signal for that specific replicate. Curves are fit with a sigmoidal equation with relevant statistics listed in Supplemental Table 2. The means ± standard deviations of 3 experimental replicates are plotted.

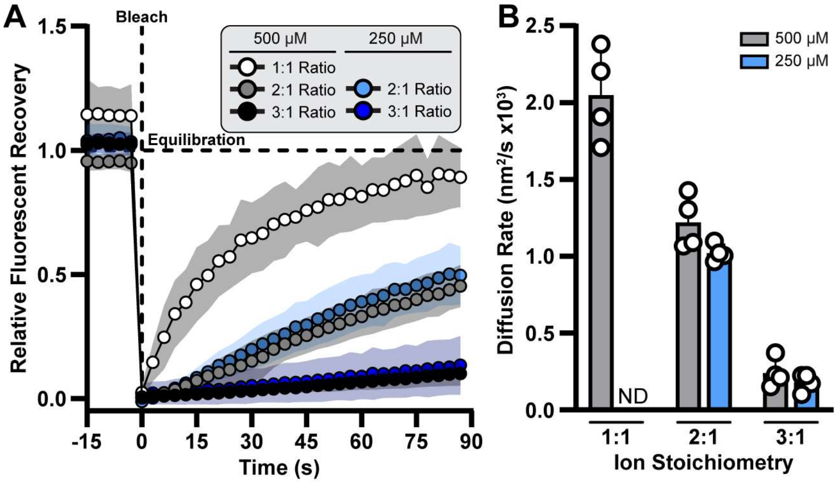


<u>**Supplemental Figure 3:**</u> **Materials related to main Figure 3. (A)** FRAP data as in main Figure 3C with the inclusion of samples prepared with 250 μM EP-LanM. Data is the mean of 4 experimental replicates ± standard deviation represented by the infill. **(B)** The diffusion rates calculated in Figure 3D with the inclusion of samples prepared with 250 μM EP-LanM. Data is the mean of 4 experimental replicates ± standard deviation. Data for the 1:1 stoichiometry at 250 μM EP-LanM could not be collected due to droplet size limitations.

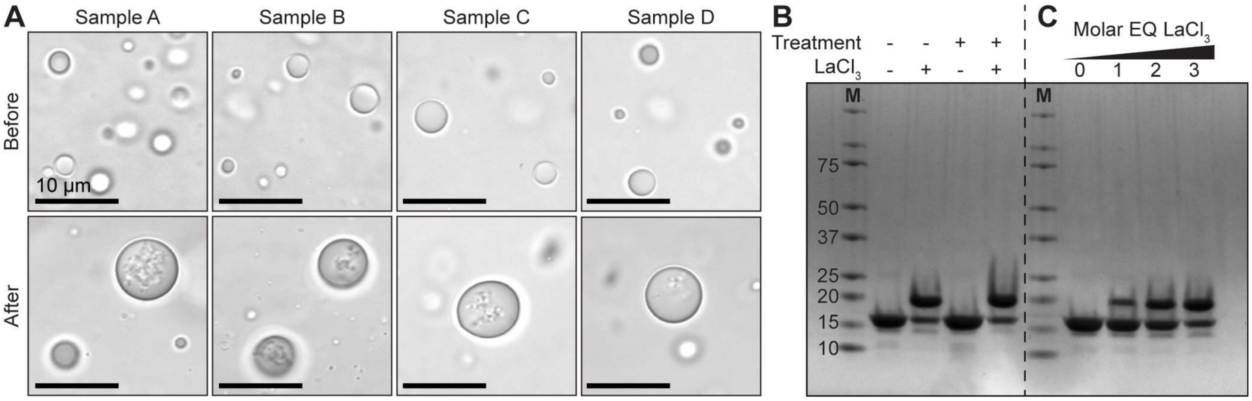


**<u>Supplemental Figure 4:</u> Materials related to main Figure 3. (A)** Light microscopy images were taken of all 4 replicates in main Figure 4C, demonstrating larger morphology and inclusions in all samples. All scale bars are 10 µm. **(B)** SDS-PAGE analysis of EP-LanM samples prepared with (+) or without (-) heat treatment or 3-molar excess of $LaCl_3$. **(C)** SDS-PAGE analysis showing the electrophoretic shift of EP-LanM in the presence of $LaCl_3$ equivalents. Both panels B and C were prepared on the same gel and cropped only to remove unused lanes. M, marker.

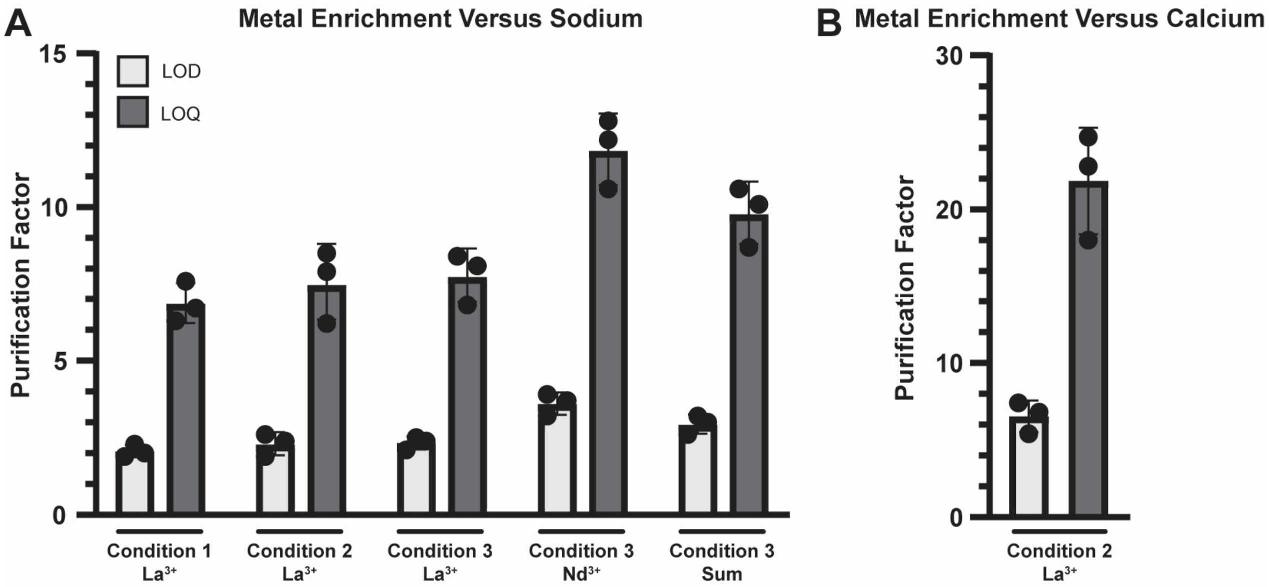


**<u>Supplemental Figure 5: Enrichment factors for metal extractions related to main Figure 5.</u>** Carry-over sodium (present in all conditions) and calcium (present only in condition 3) were detected above the limits of detection (LOD) but below the limits of quantification (LOQ). The enrichment factors were calculated for both the LOD and LOQ as these form the lower and upper bounds of metal enrichment, respectively. **(A)** The respective metal enrichment versus sodium is plotted. **(B)** The metal enrichment of $La^{3+}$ over $Ca^{2+}$ is plotted. In both panels A and B, data plotted are the means ± standard deviations of three experimental replicates.

**<u>Supplemental Table 1:</u> Sequences of LanM and EP-LanM.** This table includes the nucleotide sequences of the gene fragments used to clone LanM, EP-LanM, and mutants. The Gibson overlaps are highlighted light yellow, RBS is highlighted light blue, coding sequence highlighted light green, and N-terminal EP sequence (including linker) bolded. Mutations from wild-type are highlighted light red. Fragments were ordered from Twist Biosciences with adapters off and codon optimized for *E. coli* using the BBa_B0034 strong RBS. The PduD EP has the primary peptide sequence EINEKLLRQIIEDVLRD and the native linker has the primary peptide sequence MKGSDKPVSFNAPAASTA.

| Name | Protein | Sequence (5'->3') |
|---|---|---|
| gDT140 | LanM | TTTTTGGGCTAGCGAATTCGTACTAGAGAAAGAGGAGAAATACTAAATGGCG<br>CCAACTACGACTACCAAAGTTGATATCGCGGCGTTTGACCCGGACAAAGATG<br>GGACCATCGATCTGAAAGAGGCTTTGGCGGCAGGTTCCGCGGCCTTCGAC<br>AAGTTGGACCCGGATAAAGATGGTACTCTGGACGCCAAAGAGCTGAAGGGC<br>CGCGTGTCTGAGGCAGACCTTAAGAAGCTGGACCCTGACAATGACGGAAC<br>CCTGGACAAGAAAGAGTACTTAGCAGCGGTAGAGGCACAGTTTAAGGCCGC<br>TAACCCTGACAACGATGGCACTATTGACGCCCGTGAACTTGCAAGCCCAGC<br>GGGGTCGGCCCTGGTCAACTTAATTCGTGGTGGTGGGTCTCATCATCACCA<br>TCATCACTAAATGAGAGAAGATTTTCAGCCTGA |
| gDT141 | EP-LanM | TTTTTGGGCTAGCGAATTCGTACTAGAGAAAGAGGAGAAATACTAAATG**GAA**<br>**ATTAATGAAAAACTGCTGCGCCAGATTATTGAAGACGTACTGCGCGATATG**<br>**AAGGGCAGCGATAAACCGGTCTCGTTTAATGCGCCTGCGGCAAGCACCGC**<br>**A**GCGCCAACTACGACTACCAAAGTTGATATCGCGGCGTTTGACCCGGACAA<br>AGATGGGACCATCGATCTGAAAGAGGCTTTGGCGGCAGGTTCCGCGGCCT<br>TCGACAAGTTGGACCCGGATAAAGATGGTACTCTGGACGCCAAAGAGCTGA<br>AGGGCCGCGTGTCTGAGGCAGACCTTAAGAAGCTGGACCCTGACAATGAC<br>GGAACCCTGGACAAGAAAGAGTACTTAGCAGCGGTAGAGGCACAGTTTAAG<br>GCCGCTAACCCTGACAACGATGGCACTATTGACGCCCGTGAACTTGCAAGC<br>CCAGCGGGGTCGGCCCTGGTCAACTTAATTCGTGGTGGTGGGTCTCATCAT<br>CACCATCATCACTAAATGAGAGAAGATTTTCAGCCTGA |
| gDT186 | EP-LanM ΔEF1 | TTTTTGGGCTAGCGAATTCGTACTAGAGAAAGAGGAGAAATACTAAATG**GAA**<br>**ATTAATGAAAAACTGCTGCGCCAGATTATTGAAGACGTACTGCGCGATATG**<br>**AAGGGCAGCGATAAACCGGTCTCGTTTAATGCGCCTGCGGCAAGCACCGC**<br>**A**GCGCCAACTACGACTACCAAAGTTGATATCGCGGCGTTTGACCCGGACAA<br>AGATGGGACCATCGCGCTGAAAGAGGCTTTGGCGGCAGGTTCCGCGGCCT<br>TCGACAAGTTGGACCCGGATAAAGATGGTACTCTGGACGCCAAAGAGCTGA<br>AGGGCCGCGTGTCTGAGGCAGACCTTAAGAAGCTGGACCCTGACAATGAC<br>GGAACCCTGGACAAGAAAGAGTACTTAGCAGCGGTAGAGGCACAGTTTAAG<br>GCCGCTAACCCTGACAACGATGGCACTATTGACGCCCGTGAACTTGCAAGC<br>CCAGCGGGGTCGGCCCTGGTCAACTTAATTCGTGGTGGTGGGTCTCATCAT<br>CACCATCATCACTAAATGAGAGAAGATTTTCAGCCTGA |
| gDT187 | EP-LanM ΔEF2 | TTTTTGGGCTAGCGAATTCGTACTAGAGAAAGAGGAGAAATACTAAATG**GAA**<br>**ATTAATGAAAAACTGCTGCGCCAGATTATTGAAGACGTACTGCGCGATATG**<br>**AAGGGCAGCGATAAACCGGTCTCGTTTAATGCGCCTGCGGCAAGCACCGC**<br>**A**GCGCCAACTACGACTACCAAAGTTGATATCGCGGCGTTTGACCCGGACAA<br>AGATGGGACCATCGATCTGAAAGAGGCTTTGGCGGCAGGTTCCGCGGCCT<br>TCGACAAGTTGGACCCGGATAAAGATGGTACTCTGGCTGCCAAAGAGCTGA<br>AGGGCCGCGTGTCTGAGGCAGACCTTAAGAAGCTGGACCCTGACAATGAC<br>GGAACCCTGGACAAGAAAGAGTACTTAGCAGCGGTAGAGGCACAGTTTAAG<br>GCCGCTAACCCTGACAACGATGGCACTATTGACGCCCGTGAACTTGCAAGC<br>CCAGCGGGGTCGGCCCTGGTCAACTTAATTCGTGGTGGTGGGTCTCATCAT<br>CACCATCATCACTAAATGAGAGAAGATTTTCAGCCTGA |

**<u>Supplemental Table 2:</u> Statistics for curve fits in Figure 2A.** Curves were fit with the sigmoidal, 4PL model in GraphPad Prism version 10.6.1.

| Condition | Replicates | Top | Bottom | EC50 | Hill Slope | $R^2$ |
|---|---|---|---|---|---|---|
| 75 µM | 3 | 1.119 | 0.1047 | 2.225 | 1.074 | 0.9989 |
| 100 µM | 3 | 1.025 | 0.01143 | 2.151 | 1.293 | 0.9961 |
| 125 µM | 3 | 1.051 | 0.01089 | 2.013 | 1.300 | 0.9991 |

**<u>Supplemental Table 3:</u> Photobleaching recovery data.** Photobleaching data post-bleach in Figure 3C are fit to the equation y = A(1-exp(-x/T)) in GraphPad Prism version 10.6.1 where A is the fraction of mobile proteins (assumed to be 1), T is the recovery time, and x is the time in seconds. The diffusion rates were then calculated by the equation rate = $0.25r^2/T$ where r is the radius of the photobleached spot and T was recovery time from the FRAP profiles.

| Sample | Replicate | Recovery Time (s) | $R^2$ | Spot Radius (µm) | Diffusion Rate ($nm^2/s$) |
|---|---|---|---|---|---|
| 500 µM, 1:1 | 1 | 32.9 | 0.9976 | 0.501 | 1907.3 |
| | 2 | 13.4 | 0.9527 | 0.357 | 2380.0 |
| | 3 | 42.5 | 0.9368 | 0.612 | 2204.0 |
| | 4 | 40.4 | 0.8996 | 0.526 | 1708.4 |
| 500 µM, 2:1 | 1 | 233 | 0.9126 | 0.997 | 1066.5 |
| | 2 | 152 | 0.9880 | 0.892 | 1303.8 |
| | 3 | 106 | 0.9960 | 0.779 | 1426.8 |
| | 4 | 155 | 0.9916 | 0.821 | 1089.4 |
| 500 µM, 3:1 | 1 | 1435 | 0.9661 | 1.142 | 227.3 |
| | 2 | 500 | 0.9905 | 0.864 | 372.5 |
| | 3 | 1189 | 0.9862 | 0.848 | 151.1 |
| | 4 | 864 | 0.9843 | 0.856 | 211.9 |
| 250 µM, 2:1 | 1 | 102 | 0.9854 | 0.670 | 1098.6 |
| | 2 | 180 | 0.9743 | 0.851 | 1004.2 |
| | 3 | 174 | 0.9837 | 0.819 | 964.8 |
| | 4 | 87 | 0.9927 | 0.596 | 1020.7 |
| 250 µM, 3:1 | 1 | 1304 | 0.9000 | 1.082 | 224.2 |
| | 2 | 1136 | 0.9355 | 0.886 | 172.7 |
| | 3 | 218 | 0.7402 | 0.441 | 223.1 |
| | 4 | 1584 | 0.9600 | 0.798 | 100.4 |

**<u>Supplemental Table 4:</u> Details on EP-LanM conjugation with Alexa Fluor 488 TFP ester.** The following equations were used to quantify the protein in the labeling reaction eluant (1) and the molar ratio of dye:protein (2):

$$(1)\ \ [\text{Protein}](\text{M}) = \frac{\text{A275} - (\text{A494}\times\text{CF})}{\varepsilon} \qquad\qquad (2)\ \ \text{Dye/Protein} = \frac{\text{A494}}{\varepsilon' \text{x}[\text{Protein}]}$$

With $A_{494}$ corresponding to the absorbance at the optimal wavelength for Alexa Fluor 488 (494 nm), $A_{275}$ being the absorbance at 275 nm for LanM, CF being the correction factor (0.11) approximation for the dye, ε being the protein extinction coefficient (1400 $M^{-1}cm^{-1}$), and ε' being the extinction coefficient of the fluorescent dye at $A_{494}$ (73000 $M^{-1}cm^{-1}$). Both $A_{494}$ and $A_{275}$ were measured directly on a nanodrop and thus no dilution factor is considered.

| **Sample** | **$A_{494}$** | **$A_{275}$** | **[Protein] (µM)** | **Dye/Protein** |
|---|---|---|---|---|
| LanM | 3.738 | 0.898 | 348 | 0.147 |
| EP-LanM | 3.847 | 0.983 | 400 | 0.132 |
